# Cross-reactivity of antibodies against T cell markers in the Bank vole (*Myodes glareolus*)

**DOI:** 10.1101/2023.06.07.544031

**Authors:** Migalska Magdalena, Weglarczyk Kazimierz, Mezyk-Kopec Renata, Baliga-Klimczyk Katarzyna, Homa Joanna

## Abstract

The bank vole is a common Cricetidae rodent that is a reservoir of several zoonotic pathogens and an emerging model in eco-immunology. Here, we add to a developing immunological toolkit for this species by testing the cross-reactivity of commercially available monoclonal antibodies (mAbs) to the bank vole lymphocyte differentiation molecules and a transcription factor. We show that a combination of mAbs against CD4, CD3, and Foxp3 allows flow cytometric distinction of the main subsets of T cells: putative helper CD4+, cytotoxic CD8+ (as CD3+CD4-) and regulatory CD4+Foxp3+. We also provide a comparative analysis of amino acid sequences of CD4, CD8αβ, CD3εγδ and Fopx3 molecules for a number of commonly studied Cricetidae rodents and discuss mAb cross-reactivity patterns reported so far in this rodent family.

## 1. Introduction

The rodent family Cricetidae, a sister taxa of Muridae, is the second largest mammalian family (Steppan and Schenk, 2017; Wilson and Reeder, 2005) and taken together they form the most speciose group of mammals. By sheer numbers, wide geographic distributions and a prevalence of certain life history traits (Han et al., 2016, 2015) they include many reservoir species of zoonotic diseases, and so both groups are a subject of intense ecological, parasitological and epidemiological studies (e.g., Jonsson, Figueiredo and Vapalahti, 2010; Grzybek *et al*., 2015; Occhibove *et al*., 2022). However, a huge disparity remains in the field of immunology, where the overwhelming majority of research focusses on two species belonging to the Muridae family: mouse (*Mus musculus*) and rat (*Rattus norvegicus*). Although they are canonical model species in life sciences (including immunology), there is a growing appreciation for the potential benefits that inclusion of other species and genetically diverse populations would bring into the field (Flies, 2020). However, one major obstacle is the lack of specific immunological reagents, in particular monoclonal antibodies (mAbs), which remain essential for qualitative and quantitative description of immunological parameters. As the production of specific mAbs is generally expensive and time-consuming, cross-reactivity screens between commercially available mAbs and cell markers or cytokines of non-model species are crucial to bridge this gap.

In the present study, we focused on the bank vole, *Myodes glareolus* (Schreber, 1780), one of the most widespread Cricetidae rodents of the Palearctic (Wilson and Reeder, 2005) and a common subject of ecological, evolutionary and behavioral studies (Kotlík et al., 2022; Lonn et al., 2017; Sadowska et al., 2008; Tschirren et al., 2013). More recently, the bank vole has attracted attention as a potential model in the studies of prion diseases, as it is one of the few species readily susceptible to various prion infections (Larsen, 2016). Importantly, wild bank vole populations are known reservoirs of several zoonotic viruses, including lymphocytic choriomeningitis virus (LCMV), cowpox virus (CPXV), and Puumala virus (PUUV) (Grzybek et al., 2019) as well as spirochete *Borrelia burgdorferi* s.l., causative agent of Lyme disease in humans (Gomez-Chamorro et al., 2019). Consequently, numerous studies examined various components of both the innate and adaptive immune system of this species, such as cytokines (e.g., TNF-α), MHC molecules, and receptors (e.g., TLRs and TCRs) (Guivier et al., 2010b, 2010a; Migalska et al., 2018; Tschirren et al., 2013). At the same time, a budding immunological toolkit emerges for bank voles, with the development of recombinant cytokines (INF-gamma, Torelli et al., 2018) and permanent cell lines (Binder et al., 2019; Essbauer et al., 2011). However, to date, to the best of our knowledge, no specific mAbs directed against the main immunological markers have been developed for this species, nor systematic cross-reactivity tests have been performed.

Here, we set out to identify commercially available mAbs that would allow flow cytometric distinction of basic subsets of T cells: helper CD4+, cytotoxic CD8+ and regulatory CD4+Foxp3+ in the bank vole. To do so, we checked the cross-reactivity of several mAbs developed against CD4, CD8, CD3, and Foxp3 molecules of murid and cricetid rodents or human. Subsequently, we used a RT-qPCR assay to confirm that a combination of cross-reactive mAbs allowed identification of the CD4 and CD8-expressing T cells in this species. Finally, we provided a comparative analysis of amino acid sequences of CD4, CD8αβ, CD3εγδ and Fopx3 for a number of commonly studied Cricetidae rodents, putting into perspective mAb cross-reactivity patterns reported so far in this rodent family.

## 2. Materials and Methods

### 2.1 Antibodies

Our main aim was to identify commercially available mAbs that would specifically cross-react with bank vole T cell antigens. We focused mainly our efforts on anti-CD4 and anti-CD8 antibodies (Table 1), and further details on, e.g., manufacturers and concentrations are in Suppl. Table 1), in particular those that proved to cross-react with the antigens of relatively closely related Syrian hamster (Hammerbeck and Hooper, 2011) or were specifically developed for members of the Cricetidae family (Green et al., 2013; Rees et al., 2017). Furthermore, we tested a rat anti-human CD3 mAb (clone CD3-12 that targets an intracellular fragment of this molecule); an anti-MHC class II antibody that cross-reacted with Syrian hamster molecules (Hammerbeck and Hooper, 2011), and a highly cross-reactive mAb against Foxp3 transcription factor (Table 1).

**Table 1.**
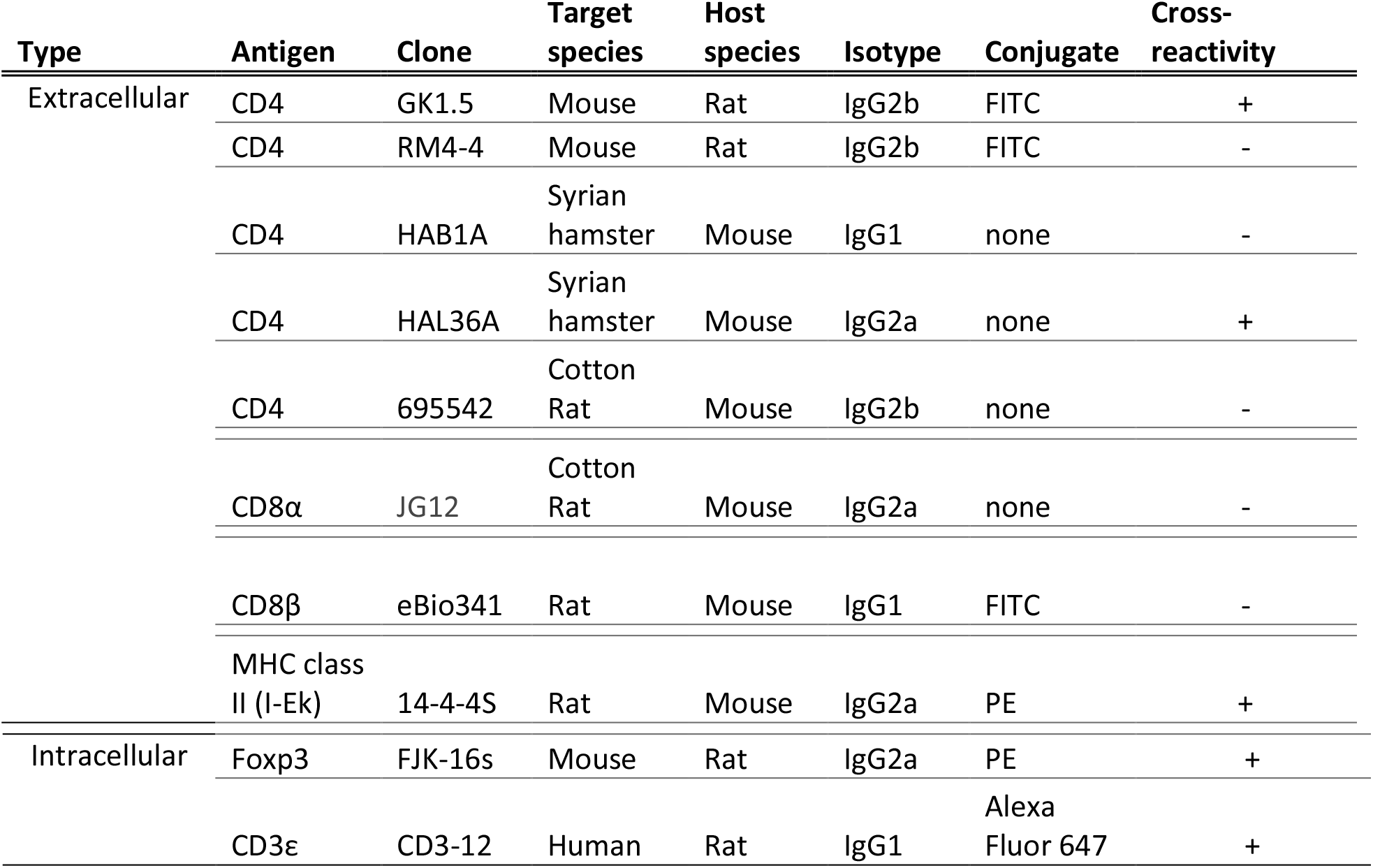
Antibodies screened for cross-reactivity in the Bank vole.

### 2.2 Animals and tissue processing

Bank voles were obtained from a laboratory colony kept at the Institute of Environmental Sciences, Jagiellonian University, Krakow (Sadowska et al., 2008), under resolution no 258/2017 of the 2nd Local Institutional Animal Care and Use Committee in Kraków. The animals were cared for according to institutional guidelines and to avoid unnecessary killing, we used animals that were removed from the colony in routine maintenance procedures that control colony size. The culled voles were of both sexes, between 2 and 7 months of age. The animals were sacrificed by cervical dislocation and spleens were harvested immediately afterwards, into high-glucose DMEM medium supplemented with 10% heat-inactivated fetal bovine serum (FBS) (Gibco, Thermo Fisher Scientific) and 1% penicillin/streptomycin. The tissues were minced and passed through a 100μm cell strainer (Biologix) into 50ml conical tubes and washed with the same medium used for the tissue harvest. The single cell suspension was centrifuged at 300-400g for 5 minutes at 12°C before resuspending in 3 ml of RBC Lysis buffer (eBioscence™) and incubated for 4 min at room temperature, with periodic gentle stroking to lyse red blood cells. Next, the cells were washed with PBS and counted.

### 2.3 Antibody cross-reactivity tests with flow cytometry

#### 2.3.1 Initial screening of cross-reactive mAbs targeting extracellular antigens

Staining was performed on 96-well U bottom plates. Cells (0.5 × 10^6^ cells/well) were blocked for 15 min in 75μl of PBS containing 2% FBS, 2% normal mouse serum (eBioscience™) and 2% normal rat serum (eBioscience™). Subsequently, antibodies targeting cell surface antigens were added in PBS containing 2% FBS, at a concentration recommended by the manufacturer (see Suppl. Table 1 for details). Cells were stained in darkness, on ice, for 30 min and washed twice with PBS. In case of unconjugated antibodies (Table 1), cells were subsequently stained with a FITC-conjugated secondary goat anti-mouse IgG (H+L) polyclonal antibody (Invitrogen, cat# 62-6511; 1:200) or PE-conjugated F(ab’)2 goat anti-mouse IgG (H+l) secondary polyclonal antibodies (eBioscience™, cat# 12-4010-82), in a buffer containing 2% FBS or 1% BSA, and stained for 30 min, in dark, on ice, and washed twice with PBS. Additionally, a portion of cells without primary antibodies was stained with secondary antibodies only, as a specificity control. Finally, cells were resuspended in PBS with 0.5% FBS for analysis on LSR Fortessa flow cytometer (BD Bioscience) equipped with CellQuest software (Becton Dickinson, San Diego, CA) or CytoFLEX (Beckman Coulter) cytometer with CytExpert Acquisition and Analysis Software (Beckman Coulter, United States).

#### 2.3.2 Extracellular and intracellular staining and cytometric analysis

After identification of cross-reactive mAbs towards extracellular antigens, we checked the cross-reactivity of mAbs targeting intracellular antigens (Table 1). Staining was performed on 96-well U bottom plates. Here, cells were first stained with either LIVE/DEAD™ Fixable Violet or Near IR (780) Viability Kit (Invitrogen, cat# L34963 or L34992, 1:1000). The staining was held in a final volume of 1ml, at RT, for 20 min, followed by a wash with 10 ml of PBS with 2% FBS. Subsequently, 0.5-1 × 10^6^ cells/well were kept for 15 min in 100μl PBS with 2% FBS to block nonspecific binding. For extracellular staining, either FITC-conjugated anti-CD4 mAb (clone GK1.5, eBioscience™, 1:100) or PE-conjugated anti-MHC class II (I-Ek) (clone 14-4-4S, eBioscience™, 1:100) was added to cells for 30 min on ice and afterwards washed twice with PBS. Next, fixation/permeabilization was performed with Foxp3/Transcription Factor Staining Buffer Set (eBioscience™) according to the manufacturer’s instructions. Briefly, cells were fixed in 200μl fixation/permeabilization working solution for 30 min at RT and then washed twice with working solution of permeabilization buffer. Washed cells were resuspended in permeabilization buffer, enriched with 2% normal mouse serum and 2% normal rat serum, and blocked for 15 minutes at RT. After blocking, PE-conjugated anti-Foxp3 mAb (clone FJK-16s, eBioscience™, 1:20) and/or Alexa Fluor 647-conjugated anti CD3 mAb (clone CD3-12, Bio-Rad, 1:100) were added directly to cells and incubated for 40 min, at RT. Appropriate fluorescence minus one (FMO) controls were prepared in the same way, and non-stained controls or cells stained only with viability dies were processed alongside. Furthermore, UltraComp eBeads™ Plus Compensation Beads (Invitrogen™) were stained according to the manufacturer’s protocol as single stained controls. After two final washes with permeabilization buffer, cells were resuspended in PBS with 0.5% FBS and analyzed either on CytoFLEX cytometer or LSRFortessa™.

Lymphocytes were gated according to forward and side scatter patterns, live cells were gated based on LIVE/DEAD™ staining, and doublets were excluded based on FSC-A vs FSC-H pattern (Supp. Figure 1). Further gating of lymphocyte populations was established using FMO and non-stained control samples.

### 2.4 Identification of CD4+ and CD8+ T cell subsets in bank vole

A combination of cross-reactive mAbs against CD3 and CD4 molecules should allow, in principle, differentiation of CD4^+^ and CD8^+^ T cell subsets, with the former identified as CD3^+^CD4^+^, and the latter as CD3^+^CD4^-^. To confirm that this is the case, we designed a qPCR assay measuring CD4 and CD8 expression in sorted populations of cell subsets. However, as our cross-reactive anti-CD3 antibody targets an intracellular part of this antigen, harsh fixation and permeabilization would damage the RNA and prevent quantification. Therefore, to preserve the quality of RNA for subsequent processing, a modification of the cell staining protocol designed for cell sorting was necessary (Channathodiyi and Houseley, 2021).

#### 2.4.1 Cell sorting and RNA extraction

The staining protocol prior cell sorting was simplified and changed based on modifications proposed by Channathodiyi & Houseley, 2021. Most importantly, fewer staining steps were performed, glyoxal (Sigma Aldrich, cat. 50649) was used to fix cells (instead of the formaldehyde-based fixative available in eBioscience™ Foxp3/Transcription Factor Staining Buffer Set), and RNase inhibitors were added to the incubation and wash solutions. Moreover, RNAse-free plasticware (including filtered tips) was used throughout and all incubation steps were held on ice. Otherwise, staining followed the protocol described above, apart from the modifications described below. For the initial extracellular staining, 5-7×10^6^ cells were resuspended in a conical tube in 200μl of PBS with 2% FBS, supplemented with 1:200 RNase Inhibitor (RI, RNasin® Plus, Promega). Cells were blocked with mouse and rat sera for 10 min and subsequently stained with FITC-conjugated anti-CD4 mAb for 30 min, as previously above.

After two washes with PBS, cells were fixed in 3% glyoxal with 20% ethanol, supplemented with 1:25 RI, in a total volume of 250μl, for 15 minutes. The glyoxal solution was prepared according to the Channathodiyi & Houseley, 2021. Subsequently, cells were washed twice with PBS supplemented with 1:150 RI and permeabilized in a working solution of a saponin-based Permeabilization Buffer (eBioscience™, cat# 00-8333), supplemented with 1:25 RI, in a total volume of 200μl, for 30 minutes. Next, mouse and rat sera were added directly to the permeabilized cells, to a final concertation of 2% each, and cells were blocked for 15 min. Following blocking, cells were stained with Alexa Fluor 647-conjugated anti CD3 mAb, for 45 min. Finally, cells were washed twice with permeabilization buffer supplemented with RI (1:100) and resuspended in PBS supplemented with RI (1:100).

The target populations were isolated using FACS Aria IIIu (Becton Dickinson). Data were acquired and analyzed using BD FACSDiva™ Software (Becton Dickinson). Lymphocytes were gated based on their FSC and SSC parameters. CD4^+^ T cells were defined as CD3^+^CD4^+^; CD8^+^ T cells were defined as CD3^+^CD4^−^; while the third population (“negative”) was defined as CD3^-^CD4^-^, and likely contained B cells, NK cells, and certain minor sub-populations of innate-like lymphocytes (Supp. Fig. 2). 100,000 cells of each population were collected in tubes with PBS supplemented with 1:100 RI. Cells were sorted to achieve purity of above 96%. Immediately after sorting, cells were centrifuged at 1800g for 3 min at 4°C, and the supernatant was removed. Cells were then lysed with 350μl of RLT buffer (Qiagen) and kept on ice until RNA extraction. RNA was extracted the same day using the RNeasy Mini Kit (Qiagen) according to the manufacturer’s instructions. The protocol included an on column DNase digestion step, with Rnase-Free DNase set (Qiagen). The RNA was eluted with RNAse-free H2O in final volume of 20μl and stored at -80°C.

#### 2.4.2 Gene expression analysis – RT-qPCR

The assay measured the expression of the *CD4* and *CD8α* genes in sorted T cell populations, as well as that of *LCK* (lymphocyte-specific protein tyrosine kinase), which was used as an additional control of T cell enrichment, since Lck is mostly expressed in T cells and is necessary for T-cell receptor signaling (Artyomov et al., 2010). *TBP* (TATA box binding protein) was chosen as a reference gene, as its expression was shown to be among the most stable in murine (Medrano et al., 2017) and bank vole (Němcová et al., 2020) spleens. The bank vole-specific target gene primers for *CD4, CD8α* and *LCK* were taken from (Migalska et al., 2019) and for the reference *TBP* gene from (Němcová et al., 2020).

Before the RT-qPCRs, 12μl RNA was reverse transcribed with Maxima H Minus First Strand cDNA Synthesis Kit (Thermo Fisher Scientific) and oligo(dT) primers, according to the manufacturer’s protocol. The calibration sample was prepared by pooling in equal proportions (before reverse transcription) RNA extracted from the three examined cell populations (i.e., CD3^+^CD4^+^; CD3^+^CD4^-^; CD3^-^CD4^-^).

RT-qPCRs were run on the Bio-Rad® CFX96 TM Real-Time PCR System (Bio-Rad) with SYBR-green based detection of PCR products. Reactions were setup using SsoAdvanced™ Universal SYBR® Green Supermix (BioRad). The final reaction volumes were 20μl, with 10 μl of Supermix, 0.5μM of each primer, and 1μl of 1.5× diluted cDNA template. The cycling conditions were: 95°C for 30s, followed by 40 cycles at 95°C for 15s and 60°C for 30s. Each sample was run in triplicate and a mean Cq (quantification cycle value) was used for the calculations. Replicates never differed more than one quantification cycle (mean SD for all assays: 0.047, range: 0.005-0.203). Non-template control for each primer pair and a duplicate of no-RT controls for each RNA isolate (with primers for the *CD8α* gene) were additionally run, and a triplicate of a calibrator sample. The melt curves of the products were inspected for each sample to confirm specific amplification. Normalized expression levels were calculated using ΔΔCt method in Bio-Rad CFX Maestro 1.1 program (v. 4.1.2433.1219).

### 2.5 Sequence analyzes

Finally, we checked whether the pattern of cross-reactivity we detect for the bank vole (or was previously reported in other cricetid rodents, Vaughn and Schountz, 2003; Hammerbeck and Hooper, 2011) correlated with sequence similarity between orthologous molecules. First, for each gene of interest (i.e., *CD3G, CD3D, CD3E, CD4, CD8A, CD8B, FOXP3*), mouse (*M. musculus*), rat (*R. norvegicus*) and human (*Homo sapiens*) high quality, annotated mRNA reference sequences were identified through the HomoloGene and/or Genbank NCBI databases. The mouse and rat sequences were then used as queries in a blastn search of a bank vole (*M. glareolus*) transcriptome (Kotlík et al., 2018).

Next, top hits were used as queries in a blastn search of the Genbank database, where deposited sequences from other Cricetidae rodents (mostly derived by automated computational analysis and annotated using the Gnomon gene prediction method) were identified. Sequences for the following common cricetid rodents were available: eastern deer mouse (*Peromyscus maniculatus*), Chinese hamster (*Cricetulus griseus*), Syrian hamster (*M. auratus*), prairie vole (*Microtus ochrogaster*), and the bank vole itself (Genbank sequence identifiers are in the Supp. Table 2). Coding sequences (CDS) were extracted from the records (based on Genbank annotation), translated into amino acid sequences, aligned with the default settings by MAFFT v. 7 (Katoh and Standley, 2013) and visualized with BioEdit (Hall, 1999). Furthermore, since anti-CD4 and anti-CD8α cotton rat-specific mAbs are commercially available, *CD4* and *CD8A* hispid cotton rat (*Sigmodon hispidus*) sequences were taken from commercially available constructs (Cotton Rat CD4 VersaClone cDNA, cat# RDC1063, R&D Systems; Cotton Rat CD8 alpha (AAL55392) VersaClone cDNA, cat# RDC0871, R&D Systems), which had been used in the construction of said antibodies. Alignments of CD sequences, with annotations of signal peptides, transmembrane regions, and ITAMs (immunoreceptor tyrosine-based activation motives) are shown in the Supp. Fig. 3, *FOXP3* alignment is in the Supp. Fig. 4.

Next, we performed a pairwise comparison of the extracellular domains (ECD) of CD molecules (as vast majority of mAbs against CDs commonly used in flow cytometry targets extracellular parts of these molecules). For Foxp3, entire CDSs were used in the comparison. The ECDs of CD molecules were extracted based on features identified above (that is, a fragment of the amino acid sequence between the signal peptide and the transmembrane domain) and a pairwise sequence identity and similarity were calculated with the MatGAT program (v2.01) (Campanella et al., 2003), using the BLOSUM62 scoring matrix for pairwise alignment.

## 3. Results

The primary focus of this article was the identification of cross-reactive antibodies to discriminate by flow cytometry the major T cell subsets in the bank vole. The test focused on four marker molecules: CD3, CD4, CD8 and Foxp3.

### 3.1 CD3

CD3 is necessary for activation and signal transduction of both CD4+ and CD8+ T cells and is considered the major marker of T cells. It is composed (in mammals) of four distinct chains, three of which (ε, γ, δ) dimerize as εγ and εδ transmembrane molecules, while the fourth chain (?) associates with a cytosolic portion of the T-cell receptor (Owen et al., 2013). Given poor cross-reactivity of antibodies targeting the extracellular part of CD3 across nonprimate mammals (Conrad et al., 2007; Hammerbeck and Hooper, 2011) we only tested an antibody against a much more conserved, cytoplasmic portion of the CD3ε molecule (Supp. Fig. 3, area shaded in grey). This antibody stained ∼ 48% of splenocytes (42 -56%, based on a sample of five, 2-3 months old, unrelated individuals from the laboratory colony) (Figure 1). Furthermore, co-staining with mAb against rat MHC class II (clone 14-4-4S) showed mutually exclusive populations (Supp. Fig. 5) – in accordance with the notion that MHC class II is not expressed on T cells (unless activated, and this phenomenon has only limited evidence from rodents; Holling, Schooten and Van Den Elsen, 2004). It is, however, expressed on professional, antigen presenting cells (Owen et al., 2013), and in the lymphocyte population analyzed herein these would most likely be B cells. Sequence identities between particular murine and cricetid CD3 chains ranged from 58-70% for CD3ε up to 69%-80% for CD3δ (Supp. Fig. 6).

**Figure 1.**
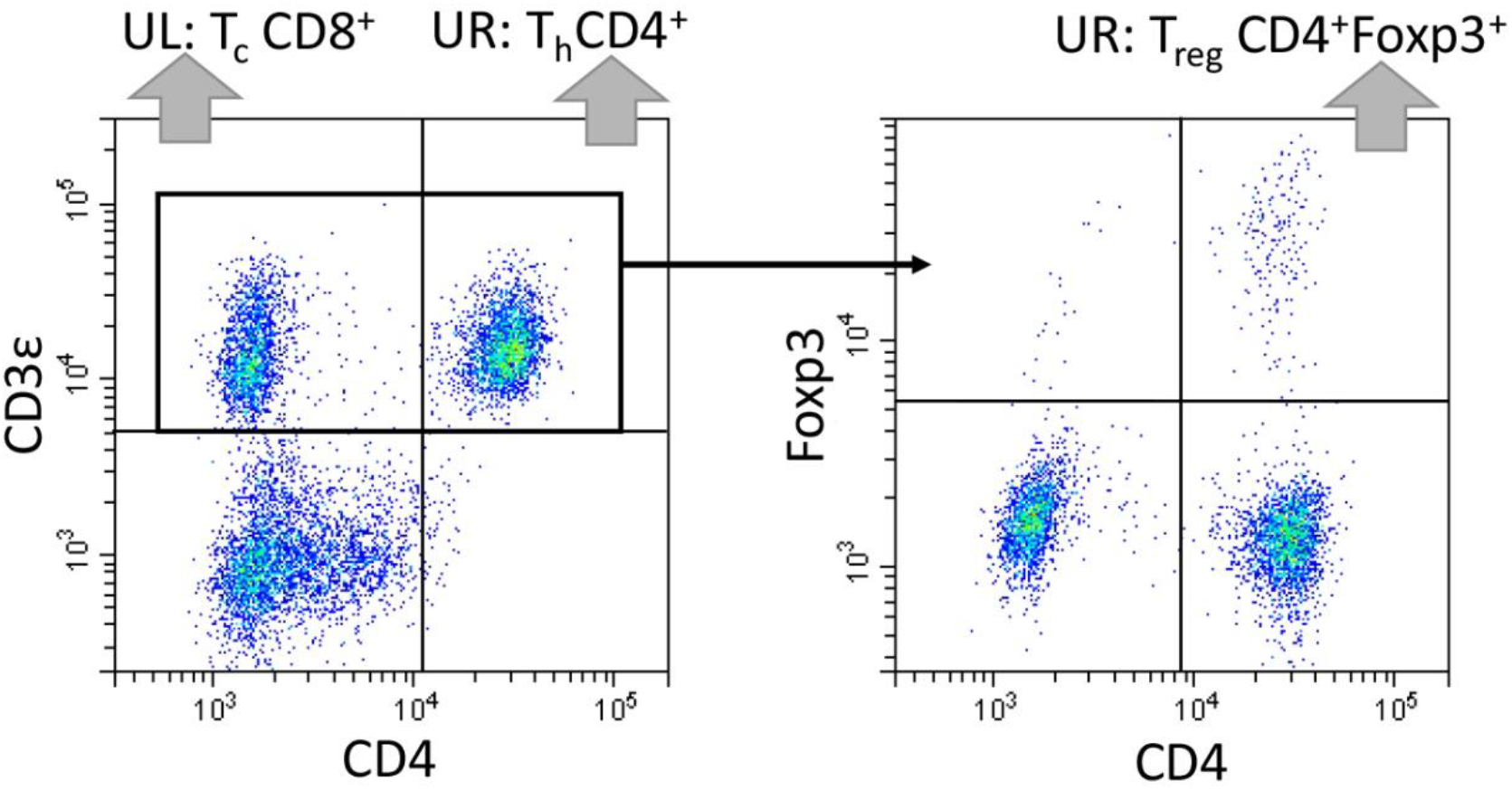
Representative Bank vole lymphocyte staining with mAbs anti-CD3 (clone CD3-12, conjugated with Alexa Fluor 647), anti-CD4 (clone GK1.5, conjugated with FITC) anti-Foxp3 (clone FJK-16s, conjugated with PE). The full gating strategy is shown in Supplementary Figure 1, details on mAbs used are in the Tab. 1 and Supp. Tab. 1.

### 3.2 CD4

CD4 is a surface monomeric glycoprotein principally expressed on T cells and thymocytes (but also on some macrophages and dendritic cells). On T cells, it serves as a co-receptor coordinating MHC class II recognition and is considered a major marker of helper T cells (Owen et al., 2013). Only two out of five tested anti-CD4 mAbs stained bank vole splenocytes – one clone developed for mouse, GK1.5 (Figure 1) and one for Syrian hamster-HAL36A. Both likely recognize the same (or similar) epitope, as suggested by the diagonal pattern of labelling reported by (Rees et al., 2017) in Syrian hamster. In contrast, another mouse-specific antibody – clone RM4-4 (which is likely to target a different epitope than GK1.5, as it does not block the binding of this mAb to CD4 – information provided by the manufacturer) did not stain bank vole cells, neither did HAB1A clone against Syrian hamster CD4 nor the 695542 clone developed for Cotton Rat CD4 (data not shown). The ECD sequence identity between bank vole and mouse CD4 was 59%, compared to 62% between the Cotton Rat and 66% between the Syrian hamster (Figure 2).

**Figure 2.**
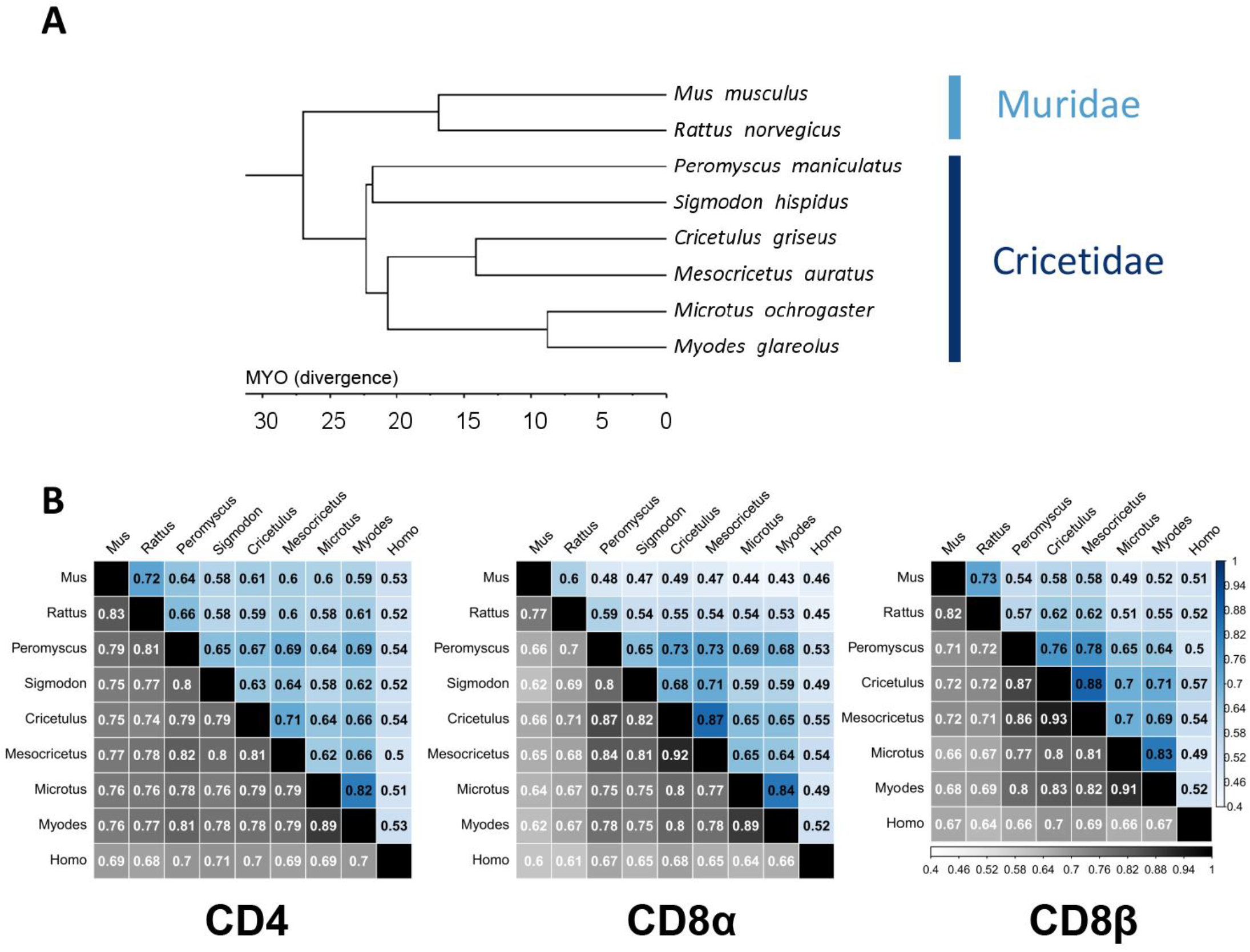
A) Tree showing phylogenetic relationships among rodent species discussed in the article. Tree topology was based on a rodent phylogenetic tree published by Steppan and Schenk (2017), with root scaled to the median divergence time between Myodes and Mus reported on Timetree (see the main text). **B) Pairwise amino acid sequence identity (upper-diagonal, in blue) and similarity (lower diagonal, in grey) matrices for CD4, CD8α and CD8β molecules**. Mus musculus – mouse, Rattus norvegicus – rat, Peromyscus maniculatus - eastern deer mouse, Sigmodon hispidus - hispid cotton rat, Cricetulus griseus - Chinese hamster, Mesocricetus auratus - Syrian hamster, Microtus ochrogaster - prairie vole, Myodes glareolus – bank vole, and Homo sapiens, human.

### 3.3 CD8

CD8 is a surface glycoprotein that can be formed by either CD8αα homodimer or CD8αβ heterodimer, and the latter form is predominantly found on cytotoxic T cells, where it serves as a co-receptor for the recognition of MHC class I (Owen et al., 2013). None of the two tested mAbs against rat CD8β or Cotton Rat CD8α chain stained bank vole cells. The sequence identity of bank vole CD8 chains compared to murid rodents was low – only 43% and 53% for CD8α (as compared to mouse and rat, respectively) and 52% - 55% for CD8β. Sequence identity with cricetid Cotton rat CD8α was only slightly higher – 59% (Figure 2).

As none of the tested, commercially available mAbs against CD8 stained bank vole cells, and low sequence similarity to murid CD8 chains gave little hope of identifying such an antibody during wider-scale testing, we checked whether a co-staining with above identified mAbs against CD3 and CD4 could be used to differentiate CD4^+^ and CD8^+^ T cell subsets (as CD3^+^CD4^+^ and CD3^+^CD4^-^, respectively).

RT-qPCR of the sorted CD3^+^CD4^+^, CD3^+^CD4^-^ and CD3^-^CD4^-^ populations confirmed high expression of both *CD4* and *LCK*, but not *CD8A* in the CD3^+^CD4^+^ cells, and high levels of *CD8*A and *LCK*, but not *CD4* in the CD3^+^CD4^-^ cells, confirming that the combination of these mAbs can be used to (at least roughly) identify the subsets of CD4^+^ and CD8^+^ T cells. Double negative cells (CD3^-^CD4^-^) showed minimal expression of *CD4*, and very low levels of *LCK* and *CD8A* (Figure 3).

**Figure 3.**
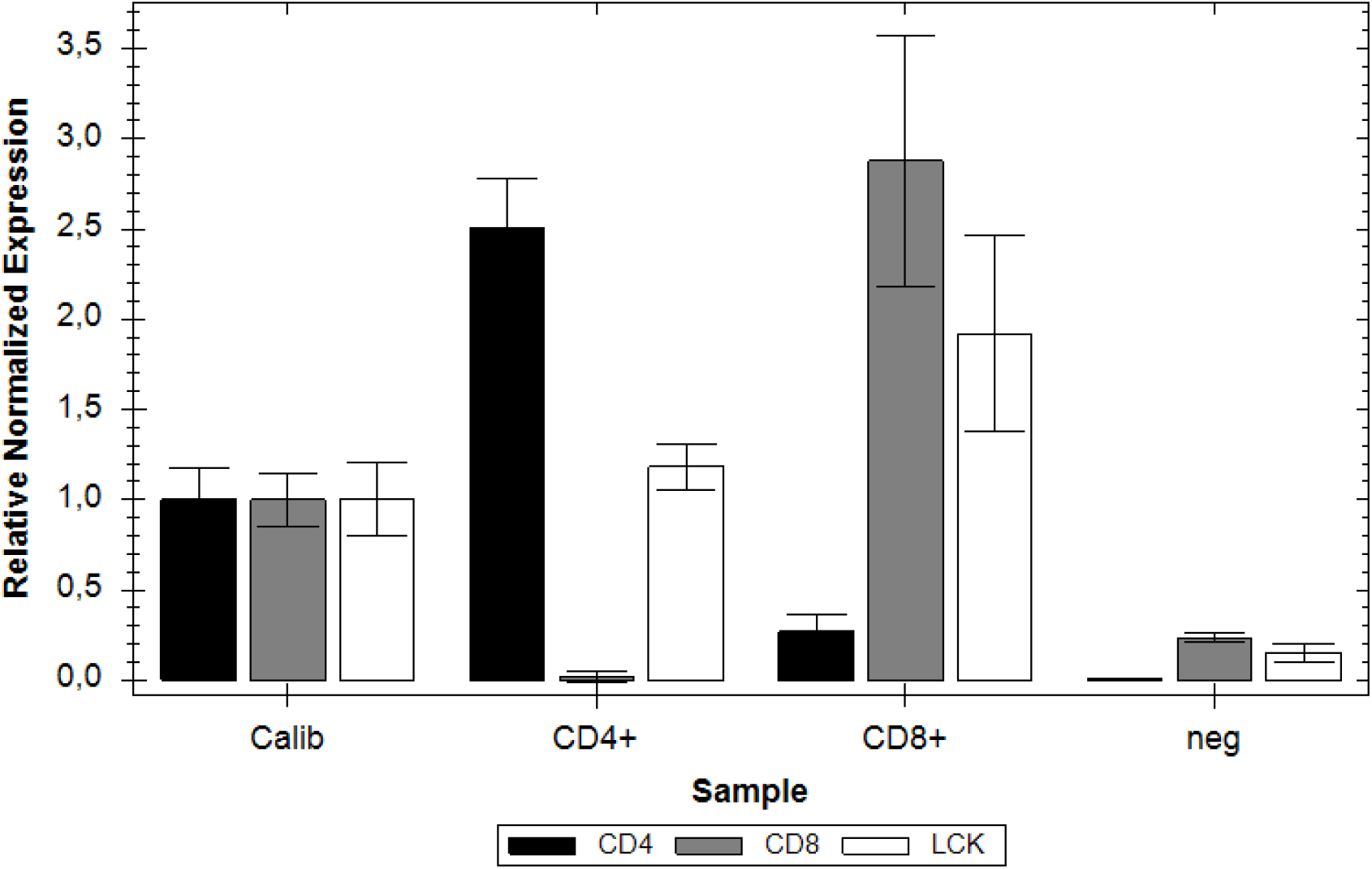
Expression of the CD4, CD8α and LCK genes in three sorted populations of cells: “CD4+” - CD3+CD4+; “CD8+” - CD3+CD4-; “neg” - CD3-CD4-. Relative normalized expression levels were measured with ΔΔCt method, with calibrator sample (“Calib”) prepared by mixing RNA from the three cell populations. TBP (TATA box binding protein) was used as a reference gene.

### 3.4 Foxp3

Foxp3 is a transcription factor that serves as a master regulator in the development and function of regulatory CD4+ T cells (Hori et al., 2017). Its sequence is highly conserved among mammals, and its identity among studied rodent species exceeded 93% (Supp. Figs. 4 and 7). In our tests, we used a highly cross-reactive rat anti-mouse Foxp3 mAb clone FJK-16s, which was proven useful in the identification of regulatory T cells in a number of mammalian species (Gerner et al., 2010; Käser et al., 2008; Rocchi et al., 2011). According to the information provided by the manufacturer, its epitope was mapped to amino acids 75-125 of the mouse Foxp3. Along this fragment, the Bank vole protein is 100% identical to the murine reference (Sup. Fig. 4, shaded in grey). On average, it stained 8% of CD4+ cells from bank vole spleens (5-10% in the individuals tested), and the staining was limited to this subset (i.e., was not found in CD3^+^CD4^-^ and CD3^-^CD4^-^ cells) (Figure 1).

## 4 Discussion

Studies devoted to the identification of cross-reactive antibodies in non-model rodents are rare, even though they could easily supplement the immunological toolbox available for many species recognized as reservoirs of zoonotic diseases. Moreover, when performed, they are rarely accompanied by a comparative sequence analysis, so it is unclear whether sequence similarity at the amino acid level can be informative when researchers strive to find a cross-reactive mAb for their species of interest. Given a limited number of such studies, there is little intuition as to whether the successful identification of cross-reactive mAb in one species is a good predictor of cross-reactivity in a close relative.

The bank vole belongs to the Cricetidae family which split from its sister taxa, Muridae (*M. musculus, R. norvegicus*), - ca. 27 MYA (TimeTree.org, accessed: 25/04/2022, (Kumar et al., 2022)) (Figure 2A), which is comparable to the divergence time between humans and macaques. Within Cricetidae itself, the earliest split divided Palearctic Cricetinae (*C. griseus, M. auratus*) and Holarctic Arvicolinae (*M. glareolus, M. ochrogaster*) from the New World clades of Neotominae (*P. maniculatus*), Sigmodontinae (*S. hispidus)*, and Tylomyinae (Steppan and Schenk, 2017).

The amino acid sequence identity of the extracellular domains of the compared rodent CD molecules was typically well below 80% (Figure 2, Supp. Fig. 4) – which is an often assumed threshold above which cross-reactivity towards orthologous molecules can be expected. Additionally, overall similarity levels differed markedly between analyzed CD molecules or even between subunits of the same molecule. When their complex, often multimeric structure, glycosylation patterns, and conformation changes upon dimerization are considered, the chances of cross-reactivity are rather slim.

CD3 is a prime example, especially given that many widely used mAbs against CD3 recognize conformational epitopes on CD3ε, expressed when bound to either CD3γ or CD3δ (Salmerón et al., 1991). This may explain why, despite comparable or higher level of sequence identity to that of CD4 molecules, finding mAbs that bind to the extracellular portion of this complex in Cricetidae can be challenging. Hammerbeck and Hooper (2011) tested nine different mAbs and were unable to identify any reactive to *M. auratus* T cells. At the same time, Vaughn and Schountz (2003) showed dimmed, but seemingly specific, staining of CD3 in *P. maniculatus* with one of these mAbs (clone 145-2C11).

The use of mAbs targeting a far more conserved cytosolic portion of CD3 molecules may thus be the most viable option in non-model rodents, as exemplified by our results in the bank vole. In general, reliance on transcription factors, such as Foxp3, and other intracellular molecules for the identification of the immune cell populations, rather than far less conserved membrane molecules, seems to be the most promising approach for non-model species. However, it should be noted that the necessary cell fixation and permeabilization of the cell membrane for intracellular staining will limit its application in, e.g. functional studies.

The patchy cross-reactivity patterns found for anti-CD4 mAbs highlight another aspect of the challenging quest that is the identification of cross-reactive antibodies. Cross-reactivity is epitope-dependent, and success in one nontarget species, while suggestive of recognition of a more conserved epitope, overall holds little predictive power for a successful staining in another, closely related species. For example, we successfully stained bank vole T cells with rat anti-mouse CD4 clone GK1.5, which also stained Syrian hamster cells (Hammerbeck and Hooper, 2011), but not those of the Cotton rat (Vaughn and Schountz, 2003). In contrast, another rat anti-mouse CD4 mAb, clone RM4-4, failed to stain cells from any of those three species.

In summary, we identified three commercially available mAbs against CD3, CD4, and Foxp3 that allow the identification of putative helper CD4+, cytotoxic CD8+ and regulatory CD4+Foxp3+ T cells in a non-model rodent form the Cricetidae family: the bank vole. We also discussed patterns of mAb cross-reactivity against these T-cell-defining molecules in the context of the levels of amino acid sequence identity among murid and cricetid rodents.

## Acknowledgements

We thank Magdalena Chadzińska and Monika Bzowska for their advice and comments on the earlier version of the manuscript. This work was funded by the National Science Centre, Poland (NSC), grant Sonatina 2019/32/C/NZ8/00440 awarded to M.M; the cost of animals was covered from the NSC grants 2016/23/B/NZ8/00888 and 2019/35/B/NZ4/03828 to Paweł Koteja.

