## Supplementary Files for "Cross-reactivity of antibodies against T cell markers in the Bank vole (*Myodes glareolus*)"

Migalska Magdalena[1], Węglarczyk Kazimierz[2], Mężyk-Kopeć Renata[3], Baliga-Klimczyk Katarzyna[1], Homa Joanna[4]

[1]Institute of Environmental Sciences, Faculty of Biology, Jagiellonian University, Gronostajowa 7, Krakow 30-387, Poland.

[2]Department of Clinical Immunology, Medical College, Jagiellonian University Department of Clinical Immunology, Institute of Paediatrics, Jagiellonian University Medical College, Wielicka 265, Krakow 30-663, Poland.

[3]Department of Cell Biochemistry, Faculty of Biochemistry, Biophysics and Biotechnology, Jagiellonian University, Gronostajowa 7, Krakow 30-387, Poland.

[4]Department of Evolutionary Immunology, Institute of Zoology and Biomedical Research, Faculty of Biology, Jagiellonian University, Gronostajowa 9, Krakow 30-387, Poland.

### List of contents

Supplementary Figure S3………………………………………………………………………………..7-13

Supplementary Figure S4……………………………………………………………………………14-15

Supplementary Figure S5……………………………………………………………………………….16

Supplementary Figure S6……………………………………………………………………………….17

Supplementary Figure S7……………………………………………………………………………….18

**Supplementary Table S1.** Antibodies screened for cross-reactivity in the Bank vole. Extension of the main text Table 1, with further details on, e.g., manufacturers and concentrations used.

| Type | Antigen | Clone | Target species | Host species | Isotype | Company | Catalogue number | Conjugate | Concentration | Used dilution |
|---|---|---|---|---|---|---|---|---|---|---|
| Extracellular | | | | | | | | | | |
| | CD4 | GK1.5 | Mouse | Rat | IgG2b | eBioscience (Thermo Fisher) | 11-0041-81 | FITC | 0.5 mg/ml | 1:100 |
| | CD4 | RM4-4 | Mouse | Rat | IgG2b | eBioscience (Thermo Fisher) | 11-0043-81 | FITC | 0.5 mg/ml | 1:100 |
| | CD4 | HAB1A | Syrian hamster | Mouse | IgG1 | Novus | NBP2-60894 | none | unknown (approx. 1mg/ml) | 1:100 |
| | CD4 | HAL36A | Syrian hamster | Mouse | IgG2a | Novus | NBP2-60895 | none | unknown (approx. 1mg/ml) | 1:100 |
| | CD4 | 695542 | Cotton Rat | Mouse | IgG2b | R&D systems | MAB6676-SP | none | unknown (approx. 0.65 mg/ml) | 1:100 |
| | CD8α | JG12 | Cotton Rat | Mouse | IgG2a | R&D systems | MAB7080-SP | none | unknown (approx. 0.7 mg/ml) | 1:100 |
| | CD8β | eBio341 | Rat | Mouse | IgG1 | eBioscience (Thermo Fisher) | 11-0080-81 | FITC | 0.5 mg/ml | 1:100 |
| | MHC class II (I-Ek) | 14-4-4S | Rat | Mouse | IgG2a | eBioscience (Thermo Fisher) | 11-0080-81 | PE | 0.2 mg/ml | 1:100 |
| Intracellular | | | | | | | | | | |
| | Foxp3 | FJK-16s | Mouse | Rat | IgG2a | Invitrogen (Thermo Fisher) | 12-5773-82 | PE | 0.2 mg/ml | 1:20 |
| | CD3ε | CD3-12 | Human | Rat | IgG1 | Bio-Rad | MCA | Alexa Fluor 647 | 0.05 mg/ml | 1:100 |

**Supplementary Table S2.** Genbank accession numbers (or other sequence identifiers) of the sequences used for comparative sequence analyses. Species used: mouse (*Mus musculus*), rat (*Rattus norvegicus*), eastern deer mouse (*Peromyscus maniculatus*), hispid cotton rat (Sigmodon hispidus), Chinese hamster (*Cricetulus griseus*), Syrian hamster (*M. auratus*), prairie vole (*Microtus ochrogaster*), bank vole (*Myodes glareolus*), human (*Homo sapiens*).

| **Species** | ***CD3G*** | ***CD3D*** | ***CD3E*** | ***CD4*** | ***CD8A*** | ***CD8B*** | ***FOXP3*** |
|---|---|---|---|---|---|---|---|
| *Mus musculus* | NP_033980.1 | NM_013487.3 | NP_031674.1 | NP_038516.1 | NM_001081110.2 | NP_033988.1 | NP_473380.1 |
| *Rattus norvegicus* | NP_001071114.1 | NM_013169.1 | NP_001101610.1; XP_008764383.1 | NP_036837.1 | NP_113726.1 | NP_113727.1 | NP_001101720.1 |
| *Peromyscus maniculatus* | XP_015863594.1 | XM_006989827.1 | XP_015863619.1 | XP_006992131.1 | XP_028716055.1 | XM_006991006.3 | XM_015988604.2 |
| *Sigmodon hispidus* | - | - | - | RDC1063* | AY065643.1 | - | - |
| *Cricetulus griseus* | XP_027267799.1 | XM_003515481.4 | XP_027267800.1 | XP_027284054.1 | XP_003503953.1 | XP_003503961.2 | XM_027433184.2 |
| *Mesocricetus auratus* | XP_005069330.1 | NM_001281420.1 | XP_021082410.1 | XP_005065990.2 | XP_005076140.1 | XP_021086297.1 | XM_005085088.4 |
| *Microtus ochrogaster* | XP_005347272.1 | XM_005347216.1 | XP_026634750.1 | XP_005365396.1 | XP_005364820.1 | XP_005364821.1 | XM_005352772.2 |
| *Myodes glareolus* | XM_048417266.1 | XM_048417267.1 | XM_048417197.1 | XM_048444203.1 | XM_048431543.1 | XM_048431869.1 | XM_048461568.1 |
| *Homo sapiens* | NP_000064.1 | NM_000732.4 | NP_000724.1 | NP_000607.1 | NP_001139345.1 | XM_011533164.3 | NP_054728.2 |

*R&D Systems™ Cotton Rat CD4 VersaClone cDNA; cat# RDC1063

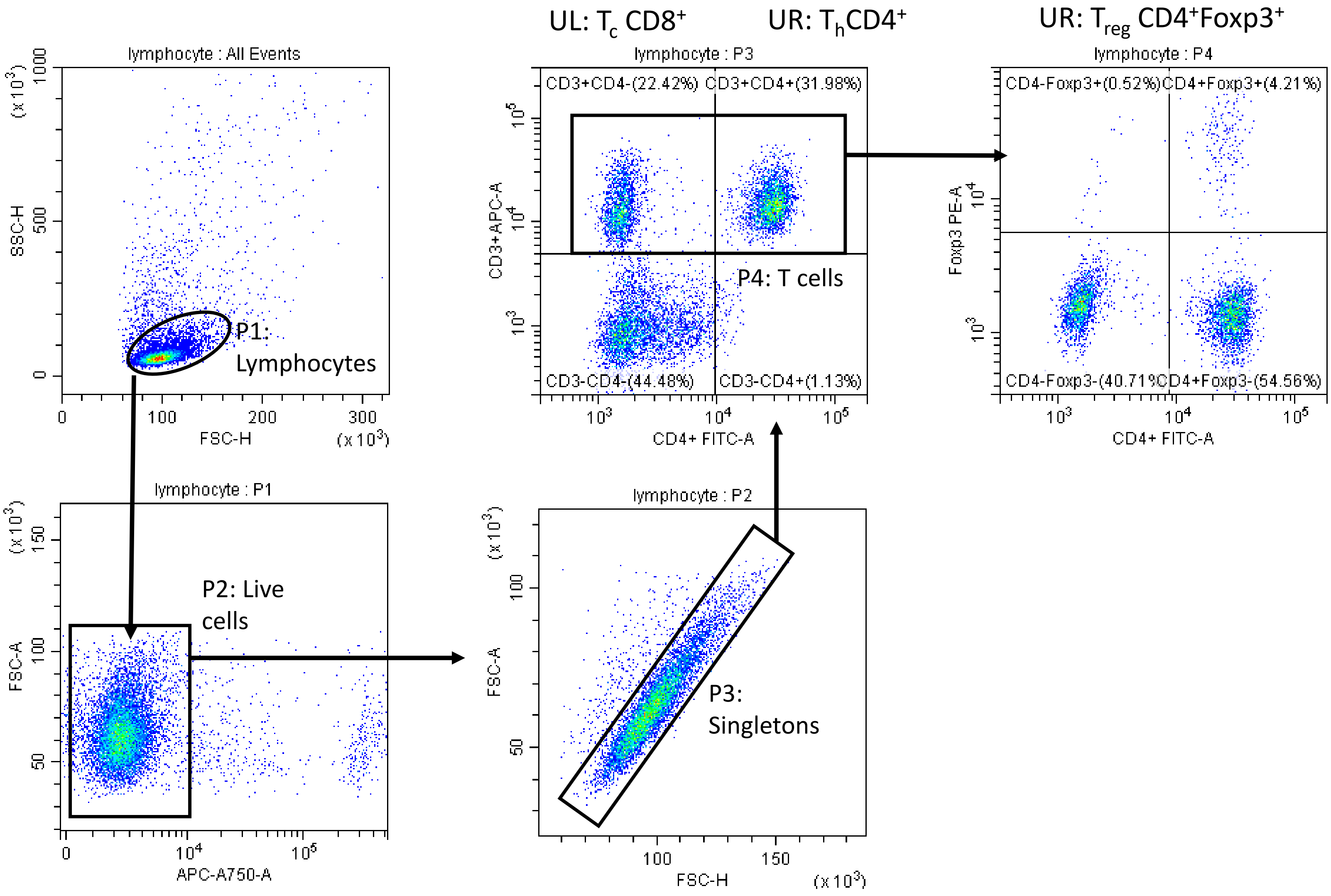

**Supplementary Figure S1.** Full gating strategy of the bank vole lymphocytes. First, lymphocytes were gated according to forward and side scatter patterns, then live cells were gated based on LIVE/DEAD™ staining, next, the doublets were excluded based on FSC-A vs FSC-H pattern. Major T cell subsets were gated based on staining with anti-CD3 (clone CD3-12, conjugated with Alexa Fluor 647), anti-CD4 (clone GK1.5, conjugated with FITC) anti-Foxp3 (clone FJK-16s, conjugated with PE).

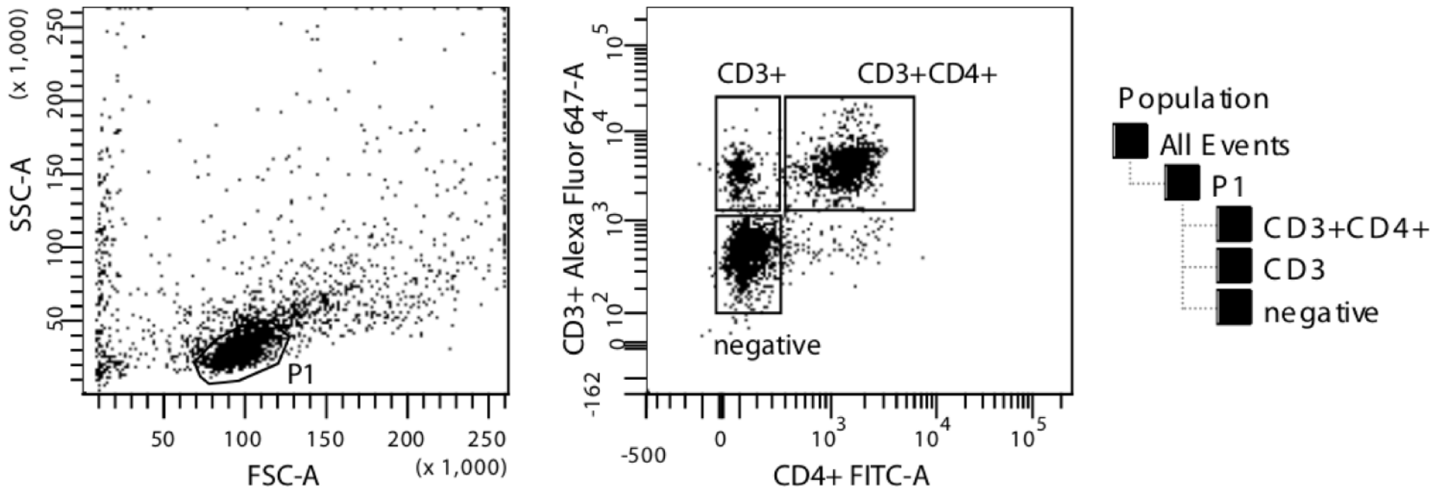

**Supplementary Figure S2.** Gating strategy for cell sorting with FACS Aria IIIu (FACSDiva™ Software, Becton Dickinson). Lymphocytes were gated based on their FSC and SSC parameters. $CD4^{+}$ T cells were defined as $CD3^{+}CD4^{+}$; $CD8^{+}$ T cells were defined as $CD3^{+}(CD4^{-})$; while the third population ("negative") was defined as $CD3^{-}CD4^{-}$, and likely contained e.g., B cells.

#### CD3 delta

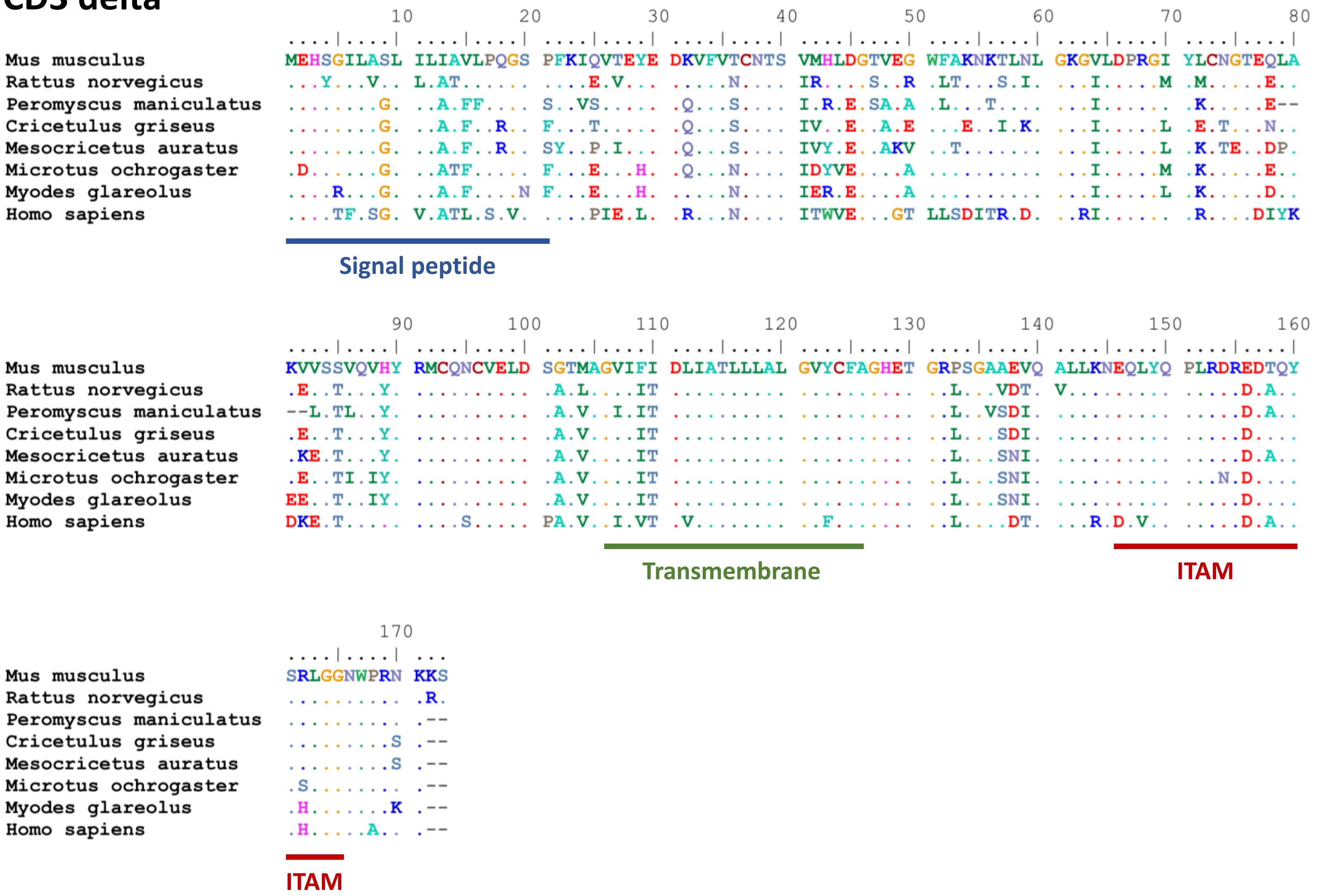

**Supplementary Figure S3.** Alignments of amino acid sequences of CD subunits present on the major T cell types (*CD3G, CD3D, CD3E, CD4, CD8A, CD8B*) with annotations of signal peptides, transmembrane regions, and ITAMs (immunoreceptor tyrosine-based activation motives). Species used: mouse (*Mus musculus*), rat (*Rattus norvegicus*), eastern deer mouse (*Peromyscus maniculatus*), Chinese hamster (*Cricetulus griseus*), Syrian hamster (*M. auratus*), prairie vole (*Microtus ochrogaster*), bank vole (*Myodes glareolus*), human (*Homo sapiens*). Additionally, for *CD8A* and *CD4* sequences from hispid cotton rat (*Sigmodon hispidus*) are shown. For *CD3E* shaded area shows region targeted by the anti-CD3 clone CD3-12 mAb.

### CD3 gamma

```
                                 10         20         30         40         50         60         70         80
                         ....|....| ....|....| ....|....| ....|....| ....|....| ....|....| ....|....| ....|....|
Mus musculus             MEQRKGLAGL FLVISLLQGT VAQTNKA--K NLVQVDGSRG DGSVLLTCGL TDKTIKWLKD GSIISPLNAT KNTWNLGNNA
Rattus norvegicus        ...G...... .......... M..QKEE--. H..K..D.Q. ........DF NE...T.... .HR...P... .S......G.
Peromyscus maniculatus   ...V...T.. V...F-.... .V.PKQENQN H..K..DN.. .D..F..... K....T.I.. .HT..S-.V. .H.....SS.
Cricetulus griseus       ..LE...T.. I...F..... ...SKQE--S H..K..DN.. ........D. K....M.... .NT..TP... .......SIT
Mesocricetus auratus     ...K...IS. I...F...-- ------E--S H.IKL.DN.. ..P.....D. K.MA.M.S.. ..T...Y.T. ........SI.
Microtus ochrogaster     ...G...T.. I...F....I ...EKQE--N H..K..DN.. ......M.D. K....T.... .NR...P... .KK....S.T
Myodes glareolus         ...G...T.. I.A.F....I ...KKQE--N H..K..DN.. ......K.D. K....T.... .N....P... ........SST
Homo sapiens             ...G....V. I.A.I..... L..SI.G--N H..K.YDYQE ........DA EA.N.T.F.. .KM.GF.TED .KK....S..
```

Signal peptide

```
                                 90        100        110        120        130        140        150        160
                         ....|....| ....|....| ....|....| ....|....| ....|....| ....|....| ....|....| ....|....|
Mus musculus             KDPRGTYQCQ GAKETSNPLQ VYYRMCENCI ELNIGTISGF IFAEVISIFF LALGVYLIAG QDGVRQSRAS DKQTLLQNEQ
Rattus norvegicus        .....M...R ...KK.QL.. ....L..... ...M..V... ....I..... ..V...F... .......... ..........
Peromyscus maniculatus   .....M.W.. .PTGK.K... .......... .......... ....I..... ..V...F... .......... ......P...
Cricetulus griseus       ......FW.. ...DK.K... .......... .......... ....I..... ..V...F... .......... ......P...
Mesocricetus auratus     .......W.. ...DK.KT.. L......... .I.....C.. ....I..... ..V....... .......... ......P...
Microtus ochrogaster     .....M.W.. ...DK.K... .......... ..SS...W.. ....I..... ..V...F... .......... ......P...
Myodes glareolus         ...Q.V.W.. ...DK.K... .......... Q.SM...... L...I..... ..V...F... .......... ......P...
Homo sapiens             .....M...K .SQNK.K... ......Q... ...AA..... L...IV...V ..V...F... .......... ......P.D.
```

Transmembrane

ITAM

```
                                170        180
                         ....|....| ....|....| ....
Mus musculus             LYQPLKDREY DQYSHLQGNQ LRKK
Rattus norvegicus        V......... E...R..... V...
Peromyscus maniculatus   ........DD .......... R...
Cricetulus griseus       ........DN .......... ....
Mesocricetus auratus     ........DD ........IP ..R.
Microtus ochrogaster     .....R..DD A.......SH P.R.
Myodes glareolus         ........DD E.......SH P.R.
Homo sapiens             .........D .......... ..RN
```

ITAM

### CD3 epsilon

```
                                 10         20         30         40         50         60         70         80
                         ....|....| ....|....| ....|....| ....|....| ....|....| ....|....| ....|....| ....|....|
Mus musculus             MRWNTFWGIL CLSLLAVGTC -QDDAENI-- ------E-YK VSISGTSVEL TCP-LDSDEN LKWEKNGQEL PQKH------
Rattus norvegicus        .Q..A..S.. G......... -.E------- ------.-.E .......... ...-.ENED. ......DKV. .D.N------
Rattus norvegicus isoX1  .Q..A..S.. G......... -.E.I.DTVT DDGARKQ-.E .......... ...-.ENED. ......DKV. .D.N------
Peromyscus maniculatus   .Q..A.CR.. L.G..L...R -...P..S-- DNEAQKY-.. ......R... ...-.ENE-V VT.K..DKTI .NEN------
Cricetulus griseus       ....A..RV. G.G..V...W -...T..P-- DNEAQKD-.T ......RI.. ...-V.GE-- IN....DKV. SEEN------
Mesocricetus auratus     ....A..RV. G....V...W -...T..P-- DNEAQKHY.. ......RI.. ...-.E..-F IH....DKV. .EEN------
Microtus ochrogaster      .Q..A..SV. G....VD..W -..ETAEN-- D-EQPKT-.. ......KI.. ...-.E.S-D I.....NK.. SGES------
Myodes glareolus         ....A..SV. G....V...L -...N.DS-- GTDAQKH-.. ......RI.. ...-I.GE-A V....DDK.. SGEI------
Homo sapiens             .QSG.H.RV. G.C..S..VW G..GN.EM-- GGITQTP-.. ......T.I. ...QYPGS-E IL.QH.DKNI GGDEDDKNIG
```

Signal peptide

```
                                 90        100        110        120        130        140        150        160
                         ....|....| ....|....| ....|....| ....|....| ....|....| ....|....| ....|....| ....|....|
Mus musculus             --DKHLVLQD FSEVEDSGYY VCYTPASN-- -KNTYLYLKA RVCEYCVEVD LTAVAIIIIV DICITLGLLM VIYYWSKNRK
Rattus norvegicus        --E.....E. ....K..... ....ES.R-- -......... ....N.M... ....S..... .......... .V.....K..
Rattus norvegicus isoX1  --E.....E. ....K..... ....ES.R-- -......... ....N.M... ....S..... .......... .V.....K..
Peromyscus maniculatus   --N.L...E. ....Q..D.. A.QAET.--- -S.A...... ....N.M... ........VI .V........ .V........
Cricetulus griseus       --..L...EN .......... A..LKD.P-- -......... ....N..... ........V. .......... .V........
Mesocricetus auratus     --E.T..... ....G..... A..LES.E-- -..A...... ....N..... .......... .......... .V.....S..
Microtus ochrogaster     --N.Q.T.N. .......... A...D.KK-- -..A...... ....N..... ........V. .......... .V........
Myodes glareolus         --E.Q.I.S. .......... T...DPKK-- -..E...... ....N..... ........V. .......... .V........
Homo sapiens             SDED..S.KE ...L.Q.... ...PRG.KPE DA.F....R. ....N.M.M. VMS..T.V.. .....G...L LV........
```

Transmembrane

```
                                170        180        190        200        210
                         ....|....| ....|....| ....|....| ....|....| ....|....| ....|
Mus musculus             AKAKPVTRGT GAGSRPR--- -GQNKERPPP VPNPDYEPIR KGQRDLYSGL NQRAV
Rattus norvegicus        .......... .T.G...GKA Q......... .......... .......... .....
Rattus norvegicus isoX1  .......... .T.G...--- -......... .......... .......... .....
Peromyscus maniculatus   .......... ..AG...--- -......... .......... .......... .....
Cricetulus griseus       .......... ...G...--- -......... .......... .......... .....
Mesocricetus auratus     ......S... ...G...--- -......... .......... .......... .....
Microtus ochrogaster     .......... ...G...--- -V.S...... .......... .A........ .....
Myodes glareolus         ........R. ...G...--- -......... .......... ....V..... .....
Homo sapiens             .........A ...G.Q.--- -......... .......... .......... ...RI
```

ITAM

# CD4

```
                                10        20        30        40        50        60        70        80
                        ....|....| ....|....| ....|....| ....|....| ....|....| ....|....| ....|....| ....|....|
Mus musculus            MCRAISLRRL L-LLLLQLSQ LLAVTQGKTL VLGKEGESAE LPCESSQKKI TVFTWKFSDQ RKILGQ--HG KGVLIRGG--
Rattus norvegicus       ...GF.F.H. .P.......K ..V......V ......G... .....TSRRS AS.A..S... KT...Y--KN .--.LIK.--
Peromyscus maniculatus  .NQG..F.H. --F.A...A. .A..S....V ......A... ....G..-.G RL.F..LP.. TR...H--QN N--FLTK.--
Sigmodon hispidus       .YQG....QF --..V...A. .P.......V KVAR..TLV. ...VG..Q.N PS.V..R... KRV.TN--LN N--.VIR.--
Cricetulus griseus      ..QG..F.H. --..V...A. .P..I....V M..R..T.I. .S.K...Q.S .F.I..LPN. TRV..M--QN N--FLIR.--
Mesocricetus auratus    .YQG..F.H. --..V...V. .P.......V L..R..K.I. ...DG..R.S A.....L... T....NNKQS N--FVARA--
Microtus ochrogaster    .NQGV.F... --..V...A. .P..L....V .V.R..NTVD ...K...E.S MF.I..L... TR...N--QN N--FVTR.RI
Myodes glareolus        .YQGV.F.H. --..V...A. .P.IL....V .V.R..DT.D ...K...E.S MF.I..L... TR...N--QN N--FVTRA--
Homo sapiens            .N.GVPF.H. --..V...AL .P.A....KV ....K.DTV. .T.TA....S IQ.H..N.N. I....N--Q. S--FLTK.--
```

Signal peptide

```
                                90       100       110       120       130       140       150       160
                        ....|....| ....|....| ....|....| ....|....| ....|....| ....|....| ....|....| ....|....|
Mus musculus            --SPSQFDRF DSKKGAWEKG SFPLIINKLK MEDSQTYICE LENRK-EEVE LWVFKVTFSP GTSLLQGQSL TLTLDSNSKV
Rattus norvegicus       --.LELYS.. ..R.N...R. .........R .......V.. ...K.-.... ....R...N. ..R....... ..I....P..
Peromyscus maniculatus  --.FEGS... ....A..DR. ....L..... ....K..... V..K.-M... ...YR..A.. D.R....... ......S-..
Sigmodon hispidus       --HFDM.N.. ..RSTL.DR. L.......VQ L...D..... VD.QR-T... ....R..AR. DYH.....T. .....CSP..
Cricetulus griseus      --QSKW.S.. ...TSE.S.. ....V.S... V...A..... V..K.MA... .......V.. D.R....... ..S...S...
Mesocricetus auratus    --QSER.S.. ..R.A..DR. ....V..... V...N..... V..K.-T... .....L.V.. DVR.....T. ..R...S...
Microtus ochrogaster    PSQSER.S.. ..R.SS.D.. ....V..... ...AGI.... V....-I... .LM.R..V.. ..H....... ....E.SL.D
Myodes glareolus        --QSER.S.. ....SS.D.. ....L..... I...GI.... V....-I... .L..R..V.. ..H....... ....E.SP..
Homo sapiens            --PSKLN..A ..RRSL.DQ. N.....KN.. I...D..... V.DQ.-...Q .L..GL.ANS D.H....... ....E.PPG-
```

```
                               170       180       190       200       210       220       230       240
                        ....|....| ....|....| ....|....| ....|....| ....|....| ....|....| ....|....| ....|....|
Mus musculus            SNPLTECKHK KGKVVSGSKV LSMSNLRVQD SDFWNCTVTL DQKKNWFGMT LSVLGFQSTA ITAYKSEGES AEFSFPLNFA
Rattus norvegicus       .D.PI..... SSNI.KD..A F.THS..I.. .GI....... N...HS.D.K ......A..S .......... ........LG
Peromyscus maniculatus  TH.SI..GGP RNSI.K...T ...P..TI.H .GI.T....Q S.Y..N.DIN I......K.S ..V..N...L .........G
Sigmodon hispidus       T..SIQ.TSP .SRLIK...T ..VP.IKT.. .GT.T....Q N.E.EK.DIN I......E.. P.V.RK...P L.V......G
Cricetulus griseus      AK.SF...GP RNG.ATH..- V.VP...I.. .GT.T.S..Q N...ETVKIN IV....KNSS TIV..K...P .......T.Q
Mesocricetus auratus    T..SMK.SGP GDR..TD.R. Y.VP...I.. .GI.T.I..Q N...ETVNID I......K.S T.V.TRD... .........G
Microtus ochrogaster    KDFSM..ASP RNGDFKS..A ITVP..SI.. HGT.S.K.A. RKQSDTLS.K I......NAS T.FF.N...A V........D
Myodes glareolus        NDLSI..TCP RKRNFKSP.A ITVP..SI.. DGT.Q....V NKQ.DTLSIK IF....KEAS T.VF.ND..L V.......LG
Homo sapiens            .S.SVQ.RSP R..NIQ.G.T ..V.Q.EL.. .GT.T...LQ N...VE.KID IV..A..KAS SIV..K...Q V......A.T
```

### CD4 continued

```
                                  250        260        270        280        290        300        310        320
                           ....|....| ....|....| ....|....| ....|....| ....|....| ....|....| ....|....| ....|....|
Mus musculus               EENG--WGEL MWKAEKDSFF QPWISFSIKN KEVSVQKSTK DLKLQLKETL PLTLKIPQVS LQFAGSGNLT LTL--DKGTL
Rattus norvegicus          ..SL--Q... R.....APSS .S..T..L.. QK.......S NP.F..S... ....Q..... .......... ...--.R.I.
Peromyscus maniculatus     ...L--R... R.R...APSP ....T..LE. .K..M..TRD N..P.ME.S. ..R....... .ES....... ...--A....
Sigmodon hispidus          D.KM--Q... K.R...VPSP EIL.T.LLE. .K..LL.TKN S.....S.A. ..H....... .EN....... .S.--T....
Cricetulus griseus         D..L--Q... ..R...A.SP G..VT..LE. .K.A.L..WG TA...M..E. ..H...H... .DN....... ...--A..K.
Mesocricetus auratus       D..L--Q... K.RT...PSP S..VT..LE. RK.TMA.D.- -R...MA.E. ..R...L... .EN....... ...--A....
Microtus ochrogaster        D..L--R... R..T..A.SP ....T..LE. .K..L..T.G N.....A... ..H....... .ES....... ...--A....
Myodes glareolus           D..L--Q... R.....APSP ....T..LE. RK..L..T.G N.....A... ..H....... .EN....... ...--A....
Homo sapiens               V.KLTGS... W.Q..RA.SS KS..T.DL.. .....KRV.Q .P...MGKK. ..H.TL..AL P.Y....... .A.EAKT.K.
```

```
                                  330        340        350        360        370        380        390        400
                           ....|....| ....|....| ....|....| ....|....| ....|....| ....|....| ....|....| ....|....|
Mus musculus               HQEVNLVVMK VAQLN-NTLT CEVMGPTSPK MRLTLKQENQ EARVSEEQKV VQVVAPETGL WQCLLSEGDK VKMDSRIQVL
Rattus norvegicus          Y......... .T.PDS.... .......... ...I...... .....RQE.. I..Q...A.V ........EE .....K....
Peromyscus maniculatus     .......... L..KD-.SV. ...R...... .K....L... D-K..RQE.. .EMKD..A.Q .L.E.N...E L.IS.K...S
Sigmodon hispidus          R......... LV.K.-.... ...R...... .....RP... RS...KQE.. .E.L...S.. .....T...E ..IN.NL...
Cricetulus griseus         R........R LI.K.-.... ...K...... .....TL... KH...KQE.. ..MP...P.. ...V.Y.AEE ..IM.DT...
Mesocricetus auratus       ....T...L. LT.N.-.I.. ...R...... .....VP.K. .....KQE.. .E.PD..A.. .R.V.Y.AEE ...NAD...S
Microtus ochrogaster       .......... L..KD-HA.I ...K....S. .Q....KD.R .VK.TK-EM. L.EP...A.. .E...K.EN. ...N.T..I.
Myodes glareolus           .......... L.MKD-.A.I ...R...... .Q....K... VD..TK-E.. F..SS..A.Q .E...N..NN ..TS.N....
Homo sapiens               .........R AT..Q-KN.. ...W...... LM.S..L..K ..K..KRE.A .W.LN..A.M ......DSGQ .LLE.N.K..
```

```
                                  410        420        430        440        450        460        470
                           ....|....| ....|....| ....|....| ....|....| ....|....| ....|....| ....|....| ..
Mus musculus               SRGVNQT--V FLACVLGGSF GFLGFLGLCI LCCVRCRHQQ RQAARMSQIK RLLSEKKTCQ CPHRMQKSHN LI
Rattus norvegicus          .K.L...--M ...V...SA. S..V.T.... .F........ .......... .......... .S........ ..
Peromyscus maniculatus     ...LK.DQPT ...L....I. S..T.I.... .......... ...E...... .......... .......TQ. ..
Sigmodon hispidus          ...WDKDQPM ...A....I. S..V.A.F.. ....K..... ...E..TH.. .......... .......TQ. ..
Cricetulus griseus         ..RL..NQPM ...V....T. S..T.V.... ..Y.K...R. ...E...... .......... .S.....TQ. F.
Mesocricetus auratus       ...L..DQPM ...I....T. S..A.M.... ....K...R. ...E...... .......... .S.....TQ. F.
Microtus ochrogaster       ...L..NQPM ..TV....T. S..A.I.... ...LK..... ...E...... .......... .......TQ. F.
Myodes glareolus           ...L..NQPM ...V....T. S..A.I.... ...IK..... ...E...... .......... .......TQ. F.
Homo sapiens               PTWSTPVQPM A.-I...VA. .L.L.I..G. FF.....RR. ...E...... .......... ....F..TCS P.
```

Transmembrane

### CD8 alpha

```
                                10        20        30        40        50        60        70        80        90
                        ....|....|....|....|....|....|....|....|....|....|....|....|....|....|....|....|....|....|
Mus musculus            MASPLTRFLSLN-LLLLGESIILGSGEAKPQAPELRIF-PKKMDAELGQKVDLVCEVLGSV-SQGCSWLFQNSSSKLPQPTFVVYMASSH
Rattus norvegicus       ...RVIC.....L...DVITRL--QV-----SGQ.QLS-...V...I..E.K.T....RDT-........R....E.L....II.VS..R
Peromyscus maniculatus  ...RV.C....T-...NAFAGL--Q.-----SKSFQMS-.PNVV..R.EN.K.I.K..L.T-.........KRGDG-LA...LI.LS.TR
Sigmodon hispidus       ..PRV....C.T-...EFIAEL--G.-----SKDFEMS-...VV.H..KE.R.T...WV.T-........LEHG.G-VK...LI.LSG.R
Cricetulus griseus      ...RV.Y....T-...DIVAEL--Q.-----LEHFQMS-...VITQ..E..E.S.Q..LTT-.........KEG.G-IK...L..VS.TR
Mesocricetus auratus    ...RVAC....T-...DVVAEL--Q.-----LEQFQMS-...VVTQ..N..E.S.Q..LTI-......I..KEG.G-VK...LI.LS.TR
Microtus ochrogaster    ...LV.C....T-...DLVAEL--Q.-----LEQ.Q.SYAA.TTP...KELK.K.KA.L.T-.........KDGPG-VK...IL.HSTAR
Myodes glareolus        ....V.C....T-...DLVAEL--Q.-----LEL.Q.S-SANTPP...KELK.K.S.RL.T-T........KDG.G-VK...LL.RS-TR
Homo sapiens            ..L.V.AL.----.P.ALLLHAAR-------PSQF.VS-.LDRTWN..ET.E.K.Q..L.NPTS.......PRGAA-AS...LL.LSQNK
```

Signal peptide

```
                               100       110       120       130       140       150       160       170       180
                        ....|....|....|....|....|....|....|....|....|....|....|....|....|....|....|....|....|....|
Mus musculus            NKITWDEKLNSSKLFSAMRDTNNKYVLTLNKFSKENEGYYFCSVISNSVMYFSSVVPVLQKVNSTTTKPVLRTPSPV-HPTGTSQ-PQRP
Rattus norvegicus       S.L--NDI.DPNL-...-.KE....I...S...TK.Q......IT........PL...F.....II....T.A.T..PP....PR-.L..
Peromyscus maniculatus  K.S--NDN.Y.NQ-I.G-EKNGDE.T...S...E..Q.......T....V...PL...FLPEKP..--..A.L.T..-PT....R-S...
Sigmodon hispidus       .ER--NN.IP.T.-L.G-KKEDK..T....N.A..D........T....V...PL.S.FLPEKP..--..PKP.TS.-PT.AI.R-SL..
Cricetulus griseus      E.R--N.D.H.I.-..G-KKNGD..T...EN.R..S........TG...VH..PL...FLPEKP..--..P.L.T..-PT....R-FV..
Mesocricetus auratus    ETR--NRD.H.IR-V.G-KKNGD..T...EN....F........TG...V...PL...FLPEKP..--..P.P.T..-AT....R-SL..
Microtus ochrogaster    I.V--ND..DENQ-..G-NKNDQE.T...K..GE..Q.......TG...V...PLL..FLPGKP..--..P.PRT..-PTV...R-SP..
Myodes glareolus        L.V--NDE.DAN.-..G-EKNG.E.T...K..GE.DQ......AT....V...PLL..FLPGKP..--.AP.PRT..-PTA...R-SP..
Homo sapiens            P.A--A.G.DTQR-..G-KRLGDTF....SD.RR.........AL...I....HF...FLPAKP...-.AP.P.T.A-PTIASQPLSL..
```

```
                               190       200       210       220       230       240       250
                        ....|....|....|....|....|....|....|....|....|....|....|....|....|....|....
Mus musculus            EDCRP--RGSVKGTGLDFACDIYIWAPLAGICVALLLSLIITLICYHRSRKRVCKCPRPLVRQEGKPRPSEKIV
Rattus norvegicus       .A...GAS...E.M..G...............AV.....V.....C..N.R..........----.......F.
Peromyscus maniculatus  .A...GAGV...KS....D.N.F.........GV.....V.....C..N.R.............G...G....F.
Sigmodon hispidus       .A...GAGT..EKK.W..D...I.L.....L.GV.....VT....C..N..........V...G...S..G.L.
Cricetulus griseus      .A...GAGV..EKR....D.N.F.........GV.....V.........N.R.............G...S....F.
Mesocricetus auratus    .A...GAGA..EKG....D.N.F.........GF...........F..N.R.........K...G...S....F.
Microtus ochrogaster    .T...GSGA..EKR....D.N.L.........GV.....V.....C..N.R.............A...S....F.
Myodes glareolus        .T...GSGA..ERR....D.N.F.......L.GV.....V.....C..N.R.............A...S..ATF.
Homo sapiens            .A...AAG.A.HTR................T.GV.....V...Y.N..N.R.........V.KSGD..SL.ARY.
```

Transmembrane

### CD8 beta

```
                                   10        20        30        40        50        60        70        80        90
                           ....|....|....|....|....|....|....|....|....|....|....|....|....|....|....|....|....|....|
Mus musculus               ----MQPWLWLVFSMKLAALWSSSALIQTPSSLLVQTNHTAKMSCEVKSISKLTSIYWLRERQDP-KDKYFEFLASWSSSKGV-LYGESV
Rattus norvegicus          ----..........V..S...G....L...........Q.......A.TFP.G.T......L..SN.N.H......RT.T..IK-...R.
Peromyscus maniculatus     MSAK..........V...........L...K..TLL..RS..VF..A..LP.T.L.H...Q.RG.A.ERR.......NP...TSV...G.
Cricetulus griseus         ----.........GV...........L...R..T.L..QV..V...A.TLP.T.L.....Q.RG.T...H.......NP...ATP...G.
Mesocricetus auratus       ----...R......V...........L...R..T.L..QI..V...A.TLP.T.V.S...Q.RG.TM..H.......NP..AAAT...G.
Microtus ochrogaster        ----..........V...........V...D.KTLL.SQALEI...ANTL..TSL.....Q.RG-S...R....V..NP....AV...G.
Myodes glareolus           ----..........V...V.......V...D.KT.L..QM.EI...ANTLPRTSL.....Q.RG-S...R....V..NP...IAV...G.
Homo sapiens               ----.R.R...LLAAQ.TV.HGN.V.Q...AYIK....KMVML...A.ISLSNMR.....Q..A.SS.SHH....L.D.A..-TIH..E.
```

Signal peptide

```
                                  100       110       120       130       140       150       160       170       180
                           ....|....|....|....|....|....|....|....|....|....|....|....|....|....|....|....|....|....|
Mus musculus               DKKRNIILESSDSRRPFLSIMNVKPEDSDFYFCATVGSPKMVFGTGTKLTVVDVLPTTAPTKKTTLKMKKKKQCPFPHPETQKGLTCSLT
Rattus norvegicus          K.NMTLSF--NS-TL...K..D......G.....M....MV..................-PT.K.T--L......T...K.......G.I
Peromyscus maniculatus     K.EDM.MS--..IS.YI.ELKS..L...GT...MMI...NLI..I..E.N..EAF..I...T.K.I--L......S.N.KA...P..G.I
Cricetulus griseus         K.ESINVF--..IT.YI.NLTS..L...GT...MM....QLI..S..E.S...A.S...Q.T.K.V--I......V.N.KA......G.I
Mesocricetus auratus       K.ENI.MF--..IT.YV.NLTS..L...GT...MMI...QLI.....E.R.......P.Q.T.K.I--L......V.N.KA......G.I
Microtus ochrogaster       K.EHM.VF--.HLS.SN.NLTRL.L...GT...MMI...QLI..K..E.R.........QPT...A--L.R.P.SVTRLKA.T.M..G.V
Myodes glareolus           K.EHI.VF--THLS.SI.NLTRL.L...GT...MMI...QLI..K............S.QPT...V--IR....RSLQLNV...T..G.V
Homo sapiens               EQEKIAVF--R.AS.FI.NLTS......GI...MI....ELT..K..Q.S...F.....QPT.KST--L..RV.RL.R......PL..PI
```

```
                                  190       200       210       220       230       240       250
                           ....|....|....|....|....|....|....|....|....|....|....|....|....|....|....|
Mus musculus               TLSLLVVCILLLLAFLGVAVYFYCVRRRARIHFMKQFHK------------------------------------
Rattus norvegicus          ......A...V..VS.S..IH.H.M..............------------------------------------
Peromyscus maniculatus     ......ASM.V..VS.S..IH.H.Q..K...R.......------------------------------------
Cricetulus griseus         ......ASL.V..VS.....HLH.L..K...........------------------------------------
Mesocricetus auratus       ......ASL.V..VS.S...HLH.L..K...R.......------------------------------------
Microtus ochrogaster       ......AS..V..VS.S...H.H.L..K...R...----------------------------------------
Myodes glareolus           ......AS..V..VS.S...H.H.L..K...R.......------------------------------------
Homo sapiens               ..G...AGV.V..VS....IHLC.R.....LR....KFNIVCLKISGFTTCCCFQILQMSREYGFGVLLQKDIGQ
```

Transmembrane

### Foxp3

```
                                  10        20        30        40        50        60        70        80        90
                         ....|....|....|....|....|....|....|....|....|....|....|....|....|....|....|....|....|....|
Mus musculus             MPNPRPAKPMAPSLALGPSPGVLPSWKTAPKGSELLGTRGSGGPFQGRDLRSGAHT-SSSLNPLPPSQLQLPTVPLVMVAPSGARLGPSP
Rattus norvegicus        ...............P.....G..................P...............S.................................
Peromyscus maniculatus   .........V.....................A........P..A.......G....--................................
Cricetulus griseus       .........V..............................P..T.......G....-.................................
Mesocricetus auratus     .........V.S.....T..........T........I..P..T.......G....--................................
Microtus ochrogaster      .........V..............................P..S.......G....--................................
Myodes glareolus         .........V..............N...............P..T.......G...A--................................
Homo sapiens             .....G...S...........AS...RA...A.D...A..P..T.......G...AS......M.........L..............L.

                                 100       110       120       130       140       150       160       170       180
                         ....|....|....|....|....|....|....|....|....|....|....|....|....|....|....|....|....|....|
Mus musculus             HLQALLQDRPHFMHQLSTVDAHAQTPVLQVRPLDNPAMISLPPPSAATGVFSLKARPGLPPGINVASLEWVSREPALLCTFPRSGTPRKD
Rattus norvegicus        ......................GH...............G....T.............................................
Peromyscus maniculatus   ..........................................................................................
Cricetulus griseus       ..........................L....................A.........................................E
Mesocricetus auratus     .........................................................................................E
Microtus ochrogaster      ..........................................................................................
Myodes glareolus         ..........................................................F...............................
Homo sapiens             .......................R......H..ES......T..TT....................................NPSA....

                                 190       200       210       220       230       240       250       260       270
                         ....|....|....|....|....|....|....|....|....|....|....|....|....|....|....|....|....|....|
Mus musculus             SNLLAAPQGSYPLLANGVCKWPGCEKVFEEPEEFLKHCQADHLLDEKGKAQCLL-QREVVQSLEQQLELEKEKLGAMQAHLAGKMALAKA
Rattus norvegicus        ..........................A....G......................-................................T..
Peromyscus maniculatus   N.............V.......................................Q................................S..
Cricetulus griseus       N.....................................................-................................S..
Mesocricetus auratus     N.....................................................-................................S..
Microtus ochrogaster      N............................Q........................-................................S..
Myodes glareolus         N.....................................................-................................S..
Homo sapiens             .T.S.V..S...................D...............R.....-...M........V......S............T..
```

**Supplementary Figure S4.** Alignment of amino acid sequences of Foxp3 transcription factor of mouse (*Mus musculus*), rat (*Rattus norvegicus*), eastern deer mouse (*Peromyscus maniculatus*), Chinese hamster (*Cricetulus griseus*), Syrian hamster (*M. auratus*), prairie vole (*Microtus ochrogaster*), bank vole (*Myodes glareolus*), human (*Homo sapiens*). Shaded area shows the region to which anti-Foxp3 mAb clone FJK-16s epitope was mapped.

### Foxp3 continued

```
                                280       290       300       310       320       330       340       350       360
                        ....|....|....|....|....|....|....|....|....|....|....|....|....|....|....|....|....|....|
Mus musculus            PSVASMDKSSCCIVATSTQGSVLPAWSAPREAPDGGLFAVRRHLWGSHGNSSFPEFFHNMDYFKYHNMRPPFTYATLIRWAILEAPERQR
Rattus norvegicus       .P...V......L..............S....S.-S...............T......................................
Peromyscus maniculatus  .AM..V..N......A.S.............PS.-S..............GT............F......................K..
Cricetulus griseus      .AM........................S....S.-S...............T............F......................K..
Mesocricetus auratus    .AM..V........TS...........S....S.-S...............T............F......................K..
Microtus ochrogaster     .AM..............A.........G....S.-S...............T............F......................K..
Myodes glareolus        .AM..V..........NA.........G....SE-S..............GT............F......................K..
Homo sapiens            S....S..G......AGS..P.V....G......-S...............T....L.......F......................K..

                                370       380       390       400       410       420       430
                        ....|....|....|....|....|....|....|....|....|....|....|....|....|....|...
Mus musculus            TLNEIYHWFTRMFAYFRNHPATWKNAIRHNLSLHKCFVRVESEKGAVWTVDEFEFRKKRSQRPNKCSNPCP--
Rattus norvegicus       ...............................................................S.......--
Peromyscus maniculatus  ..........C....................................................S.......--
Cricetulus griseus      ...............................................................S.......--
Mesocricetus auratus    ...............................................................S.......--
Microtus ochrogaster     ...............................................................S.......--
Myodes glareolus        ...............................................................S.......--
Homo sapiens            ..............F.....................................L..........SR....T.GP
```

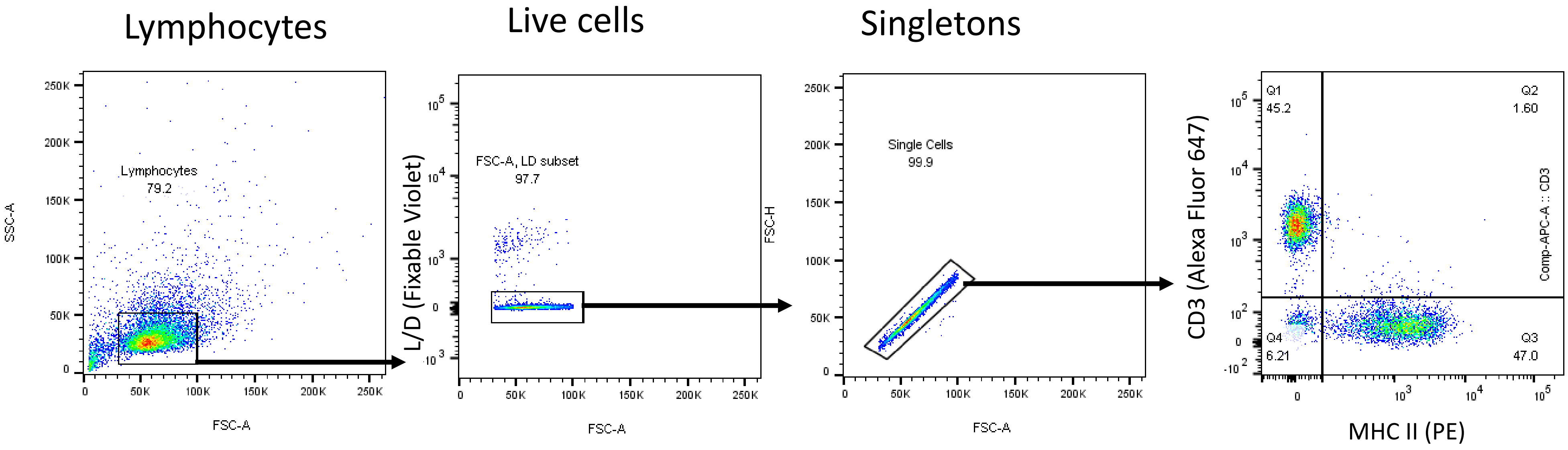

**Supplementary Figure S5.** Bank vole lymphocyte staining with anti-CD3 mAb (clone CD3-12, Alexa Fluor 647-conjugated) and with mAb against rat MHC class II (clone 14-4-4S, PE-conjugated).

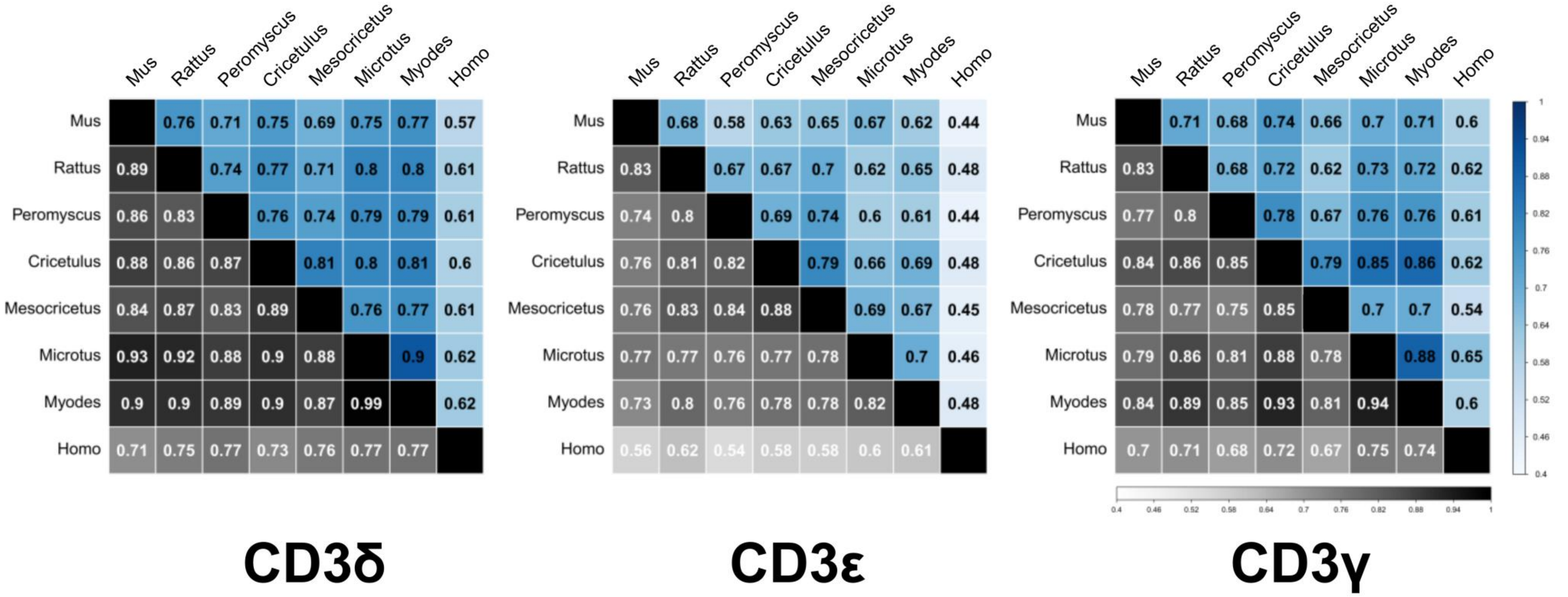

**Supplementary Figure S6.** Pairwise amino acid sequence identity (upper-diagonal, in blue) and similarity (lower diagonal, in grey) matrices for *CD3G, CD3D, CD3E* molecules. *Mus musculus* – mouse, *Rattus norvegicus* – rat, *Peromyscus maniculatus* - eastern deer mouse, *Cricetulus griseus* - Chinese hamster, *Mesocricetus auratus* - Syrian hamster, *Microtus ochrogaster* - prairie vole, *Myodes glareolus* – bank vole, and *Homo sapiens*, human.

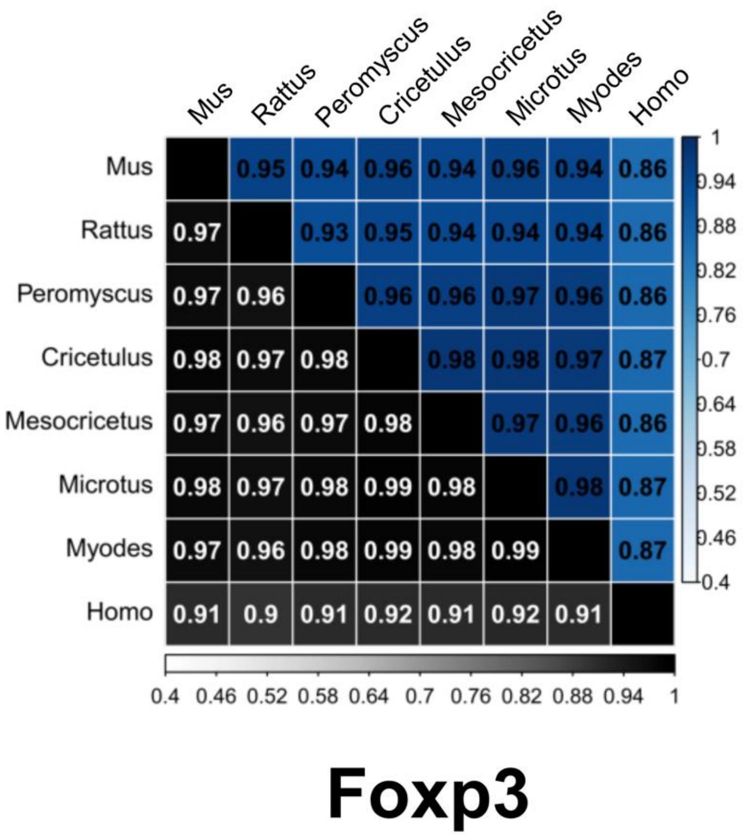

**Supplementary Figure S7.** Pairwise amino acid sequence identity (upper-diagonal, in blue) and similarity (lower diagonal, in grey) matrices for *FOXP3* molecules. *Mus musculus* – mouse, *Rattus norvegicus* – rat, *Peromyscus maniculatus* - eastern deer mouse, *Cricetulus griseus* - Chinese hamster, *Mesocricetus auratus* - Syrian hamster, *Microtus ochrogaster* - prairie vole, *Myodes glareolus* – bank vole, and *Homo sapiens*, human.
